# Accelerated loss of hypoxia response in zebrafish with familial Alzheimer’s disease-like mutation of Presenilin 1

**DOI:** 10.1101/526277

**Authors:** Morgan Newman, Hani Moussavi Nik, Greg T. Sutherland, Nhi Hin, Woojin S. Kim, Glenda M. Halliday, Suman Jayadev, Carole Smith, Angela Laird, Caitlin Lucas, Thaksaon Kittipassorn, Dan J. Peet, Michael Lardelli

## Abstract

Ageing is the major risk factor for Alzheimer’s disease (AD), a condition involving brain hypoxia. The majority of early onset familial AD (EOfAD) cases involve dominant mutations in the gene *PSEN1. PSEN1* null mutations do not cause EOfAD. We exploited putative hypomorphic and EOfAD-like mutations in the zebrafish *psen1* gene to explore the effects of age and genotype on brain responses to acute hypoxia. Both mutations accelerate age-dependent changes in hypoxia-sensitive gene expression supporting that ageing is necessary, but insufficient, for AD occurrence. Curiously, the responses to acute hypoxia become inverted in extremely aged fish. This is associated with an apparent inability to upregulate glycolysis. Wild type *PSEN1* allele expression is reduced in post-mortem brains of human EOfAD mutation carriers (and extremely aged fish), possibly contributing to EOfAD pathogenesis. We also observed that age-dependent loss of HIF1 stabilisation under hypoxia is a phenomenon conserved across vertebrate classes.

## Introduction

Alzheimer’s disease (AD) is the most prevalent form of dementia. Decreased levels of soluble amyloid-beta (Aβ) peptides in cerebrospinal fluid [1] (presumably due to decreased clearance from the brain) is regarded as one of the earliest markers of both early-onset familial AD (EOfAD) and late-onset sporadic AD (LOsAD), preceding disease onset by 20-30 years [2, 3], while vascular changes may occur even earlier [4]. When AD clinical symptoms eventually become overt, neurodegeneration may be too advanced for effective therapeutic intervention. This may explain the failure of drugs designed to inhibit supposed Aβ-related pathological processes to halt cognitive decline [5, 6].

There are numerous observations implicating brain hypoxia in AD. For example, cardiovascular risk factors strongly correlate with LOsAD [7, 8]. Other significant LOsAD risk factors include vascular brain injury and traumatic brain injury [9], while recent studies suggest that aerobic exercise can reduce AD risk [10, 11]. Reduced blood flow has been observed in AD brains (reviewed by [12, 13]) most likely causing impaired oxygenation. Hypoxia can induce the expression of EOfAD genes/proteins [14-17] to drive Aβ production and a serum biomarker associated with hypoxia can predict which people with Mild Cognitive Impairment (MCI) will progress to AD and which will not [18]. An important controller of cellular responses to hypoxia is the heterodimeric transcription factor HYPOXIA INDUCIBLE FACTOR 1, HIF1 which is comprised of HIF1α and HIF1β monomers. HIF1α protein is decreased in LOsAD brains compared to age-matched controls [19], although the reason for this is unknown.

The molecular events underlying LOsAD are complex and have eluded detailed understanding. However, EOfAD is caused by inherited, dominant mutations in the genes, *PRESENILIN 1* (*PSEN1*), *PRESENILIN2* (*PSEN2*), *AMYLOID BETA A4 PRECURSOR PROTEIN* (*APP*) and *SORTILIN-RELATED RECEPTOR* (*SORL1*) [20, 21] and so is amenable to genetic analysis in animal models. Only 5.5% of AD is defined as EOfAD [22]. Nevertheless, the similarity in clinical disease progression and brain pathology of EOfAD and LOsAD [23] supports the assumption that genetic analysis of EOfAD will contribute to understanding of both forms of the disease.

PSEN1 and PSEN2 are alternative catalytic components of the γ-secretase complexes that cleave APP to form Aβ. On a population basis, around 60% of all EOfAD mutations occur in *PSEN1* [24] and are viewed mostly as altering the cleavage of APP to yield changes in the relative abundance of different Aβ size forms. Nevertheless, the exact relationship between EOfAD mutations in the *PRESENILIN* genes and *APP* is still a matter of debate [25, 26].

Structural and functional brain imaging (MRI and/or PET) from individuals with EOfAD mutations has revealed changes as early as nine years of age [27, 28]. Elevations in Aβ deposition and reduced glucose metabolism have also been observed in EOfAD brains years before symptom onset [29]. Surprisingly, there has been little detailed molecular investigation of the young adult brains of any animal model closely mimicking the human EOfAD genetic state – i.e. heterozygous for an EOfAD-related mutation in a single, endogenous gene. Therefore, to model and explore early changes in the brain driving AD pathogenesis, we have edited the zebrafish genome to introduce EOfAD-related mutations into the endogenous, *PSEN1*-equivalent gene of zebrafish, *psen1*. We recently published transcriptomic analyses of two such mutations, *psen1*^*K97fs*^ and *psen1*^*Q96_*K97del^ [30, 31].

Analysis of the *psen1*^*K97fs*^ mutation in young and aged brains revealed that this mutation accelerates elements of brain aging [31]. However, we now know that this frameshift mutation differs from its human archetype (*PSEN2*^*K115fs*^) in failing to restore the open reading frame (ORF) of various frameshifting alternative transcripts [31, 32]. (*All* EOfAD alleles of *PSEN1* and *PSEN2* follow the “reading frame preservation rule” by producing transcripts encoding altered protein sequences without premature termination codons [25, 33].) *psen1*^*K97fs*^ most likely represents a protein-truncating, hypomorphic allele similar to the human *PSEN1*^*P242fs*^ allele that causes the skin disease familial hidradenitis suppurativa without EOfAD [34]. In contrast, the more EOfAD-like *psen1*^*Q96_K97del*^ mutation precisely deletes two codons thus following the reading frame preservation rule. It resembles numerous human EOfAD mutations by altering the structure of the first lumenal loop of PSEN1 (a multipass transmembrane protein of the endoplasmic reticulum and other organelles, [35]).

Gene ontology analysis of young adult zebrafish brain transcriptomes revealed that, in contrast to *psen1*^*K97fs*^, the more EOfAD-like *psen1*^*Q96_K97del*^ mutation is predicted to have very significant effects on mitochondrial function, especially synthesis of ATP, and on ATP-dependent functions such as the acidification of lysosomes that are critical for autophagy [36]. The importance of oxygen for production of ATP by mitochondria, the apparent role of hypoxia in AD, and the little-studied interaction between PSEN1 protein and HIF1α [37, 38] led us to question whether our two zebrafish *psen1* mutants would differ in their responses to hypoxia with age.

We found that zebrafish brain gene expression responses to hypoxia increase greatly with age but that extremely aged brains “invert” to show opposite responses. The hypomorphic and the EOfAD-like mutations of *psen1* both accelerate the onset of this inversion. Expression of the *psen1* gene itself under normoxia also increases with age until this reverses in extremely aged brains. Loss of *PSEN1* expression is also observed in human EOfAD brain tissue compared to age-matched controls supporting that these brains are subject to accelerated aging. The more EOfAD-like mutation of *psen1* causes an abnormal, and possibly pathogenic, stabilisation of a zebrafish HIF1α paralogue that becomes evident in aged brains.

## Results

The ability to generate large families of siblings (over 100 individuals) and hold them in the same environment (the same tank) reduces genetic and environmental noise in -omics analyses while allowing sampling of large numbers of individuals for statistical power. We have previously exploited this in our studies of EOfAD-related mutation model brain transcriptomes [31, 32]. In these studies, we typically generate families in which half the siblings are heterozygous mutants (modelling the human carriers of dominant EOfAD mutations) and half the siblings are non-mutant (i.e. wild type). Serendipitously, not all the individuals of these large families are required for the -omics analyses, so that follow-up experiments can be performed on individuals of various ages from families generated at various times. To investigate the interaction between age and our mutations on responses to hypoxia we sampled families extant in our zebrafish facility. As the individuals involved were those surplus to the needs of planned -omics studies, it was not always possible to analyse equal numbers of females and males. However, principal component analysis (PCA) of transcriptome data has not revealed sex to be a major determinant of zebrafish brain molecular state (see Supplementary Figure 1).

We exposed zebrafish to acute hypoxia by placement in oxygen-depleted water (0.6 ± 0.2 mg/L O_2_) for 3 hours after which they were rapidly euthanised and their brains removed for analysis.

### Accelerated loss of hypoxia responses in aging mutant zebrafish brains

During hypoxia, cells promote survival by adjusting their energy metabolism. In part, this is controlled at the transcriptional level by changes in the expression of hypoxia response genes, (HRGs, reviewed by [39]). These genes include *PDK1* encoding PYRUVATE DEHYDROGENASE KINASE1 that phosphorylates PYRUVATE DEHYDROGENASE to inhibit conversion of pyruvate to acetyl-CoA [40]. In this way, pyruvate is directed away from the tricarboxylic acid cycle (TCA) and oxidative phosphorylation and into ATP production via anaerobic glycolysis. Thus, *PDK1* acts as a form of “rheostat” to set the relative rates of anaerobic glycolysis and oxidative phosphorylation. To determine the effects of age and our *psen1* alleles on genetic responses to hypoxia, we used digital quantitative PCR (dqPCR) to assess transcript levels from four HRGs: *cd44a, igfbp3, mmp2* (reviewed in [41]) and *pdk1* [42] in the brains of zebrafish exposed to normoxia or acute hypoxia.

Unexpectedly, the responses of heterozygous zebrafish carrying either of our two *psen1* mutant alleles were essentially identical and very different to those of wild type siblings (Figure 1A, B, Tables 1 and 2, Supplementary Data 1). Young, 6-month-old mutants show increased basal levels of HRG expression under normoxia suggestive of pre-existing hypoxic stress, although other explanations are possible such as alteration of iron homeostasis or alteration of basal γ-secretase levels (see the Discussion). Nevertheless, both wild type and mutant young adult zebrafish were able to increase HRG expression under acute hypoxia. Aged, 24-month-old wild type zebrafish showed increased basal HRG expression under normoxia, resembling HRG expression in the young mutant animals (Figure 1A, B, Tables 1 and 2, Supplementary Data 1). This is consistent with our previous observation that the *psen1*^*K97fs*^ mutation accelerates brain aging [31]. Despite their increased level of HRG expression under normoxia, the 24-month-old wild type brains could still respond further to hypoxia with an approximately two-fold increase in HRG expression. Interestingly, their levels of HRG expression were then very similar to those of normoxic 24-month-old *psen1*^*K97fs*^ and *psen1*^*Q96_K97del*^ mutant zebrafish brains (Figure 1A, B, Tables 1 and 2, Supplementary Data 1), again supporting that these mutant brains are under hypoxic stress.

**Table 1.**
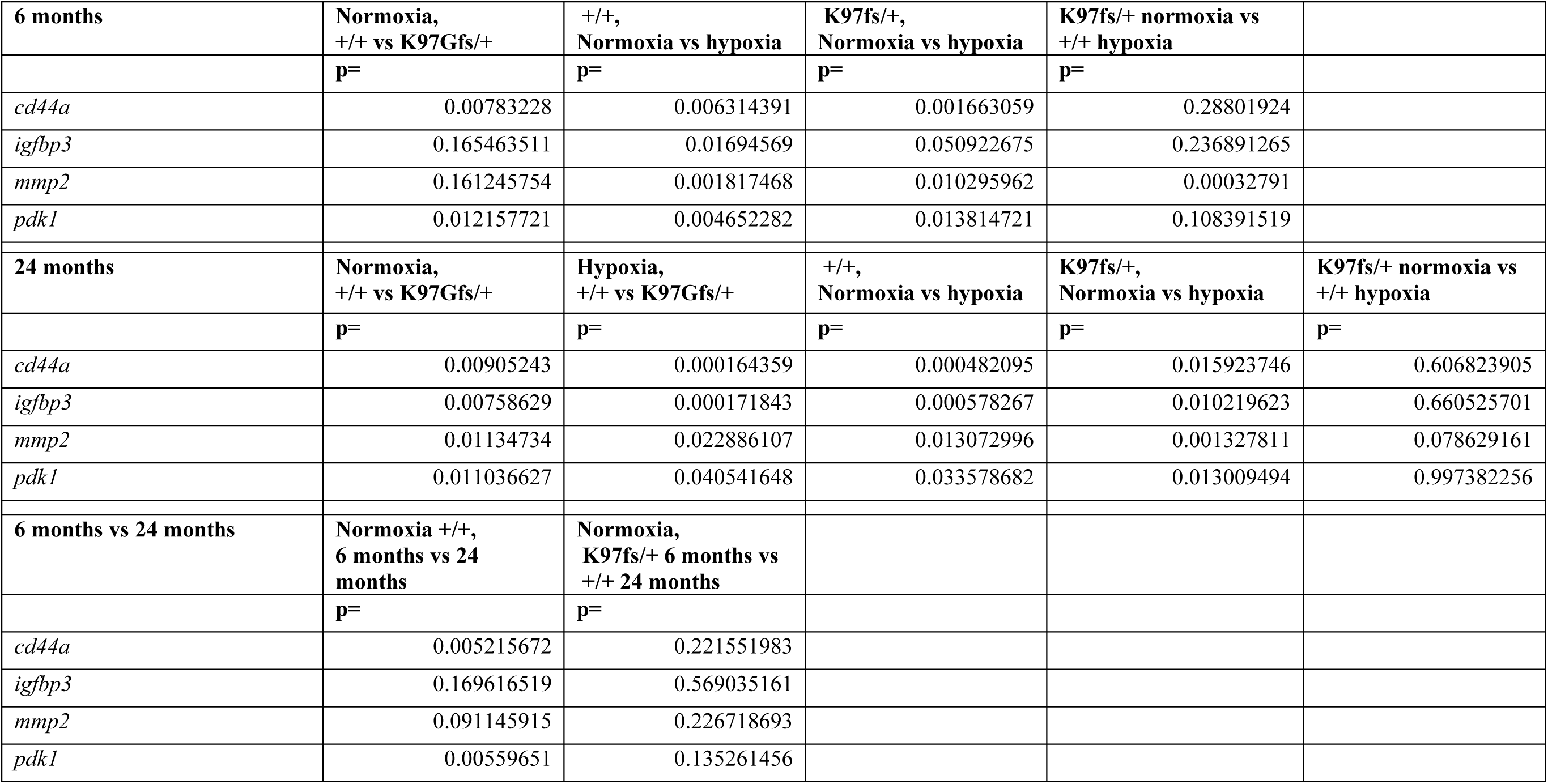
p-values calculated for comparisons of hypoxia response gene expression in 6-month-old and 24-month-old zebrafish brains under normoxia and hypoxia (K97fs/+ vs +/+ siblings).

**Table 2.**
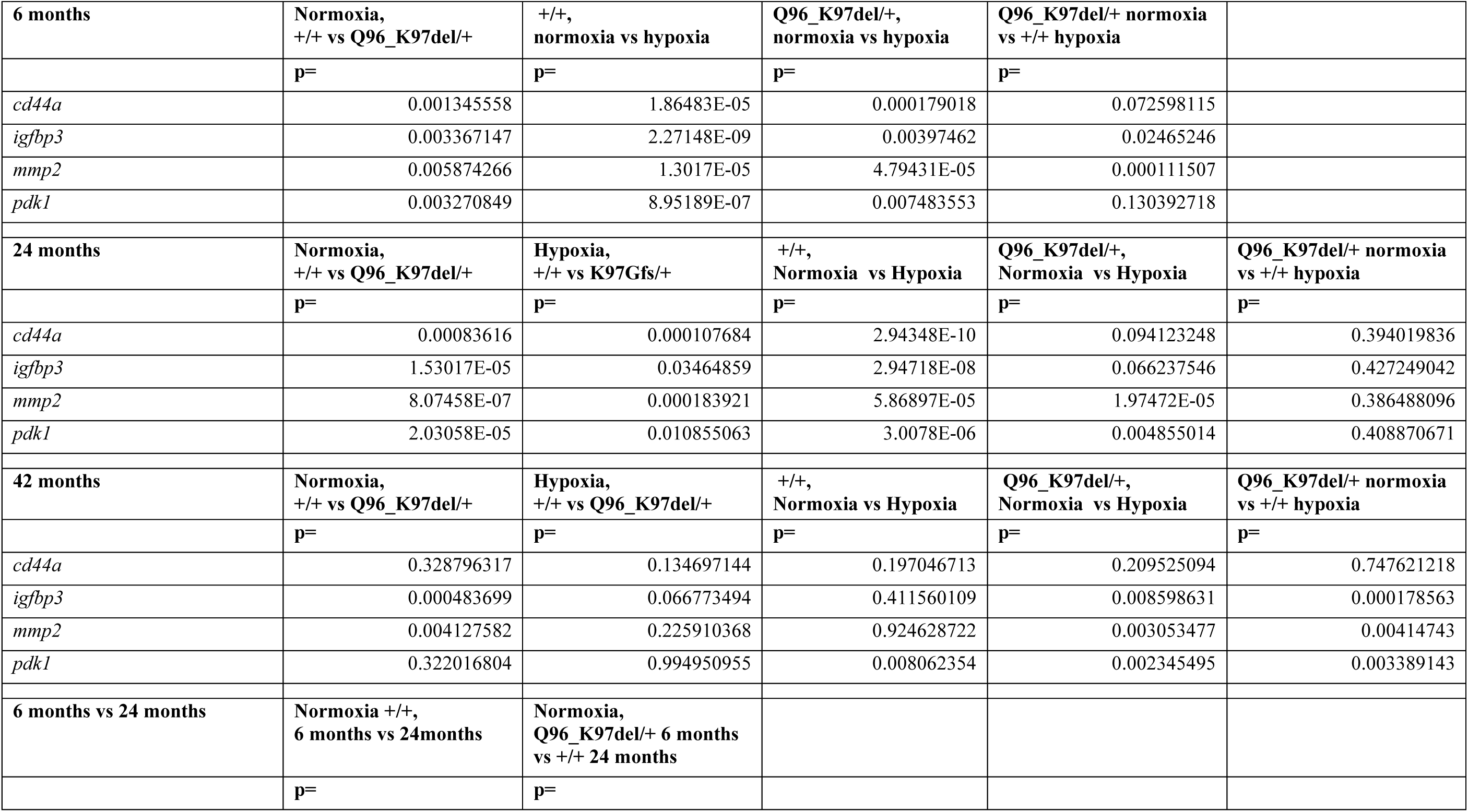
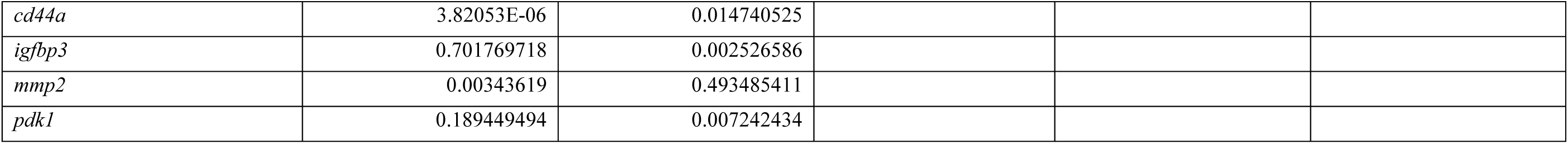
p-values calculated for comparisons of hypoxia response gene expression in 6-month-old and 24-month-old zebrafish brains under normoxia and hypoxia (Q96_K97del/+ and +/+ siblings).

**Table 3.**
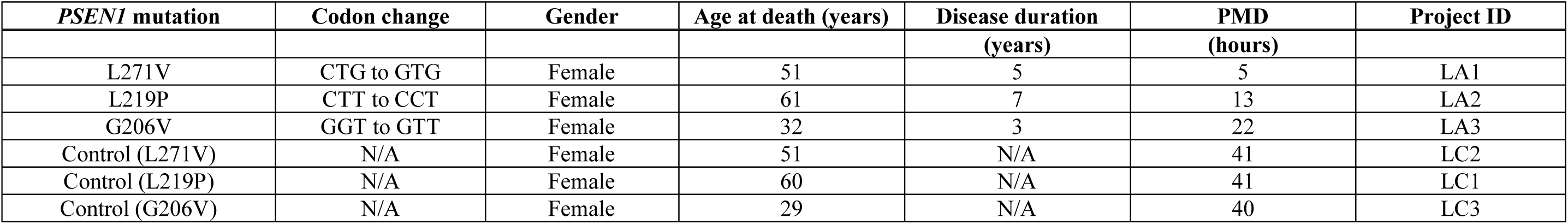
Human *PSEN1* mutations analysed for allele-specific dqPCR including age-matched control details. Post-mortem delay (PMD).

**Table 4.**
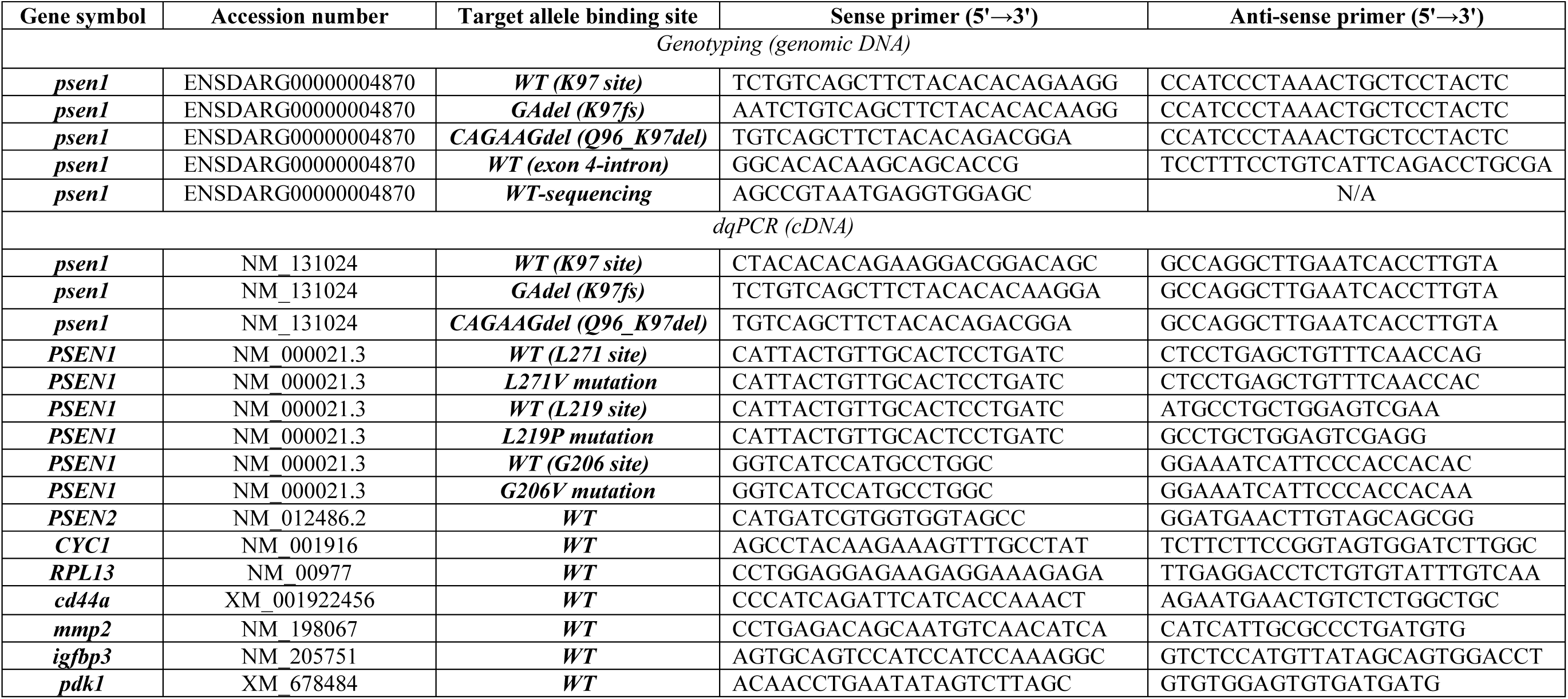
Primer sequences for genotyping, sequencing and dqPCR. WT denotes wild type allele-specific PCR. The WT primer target site for *psen1* or *PSEN1* is indicated in parentheses.

**Figure 1.**
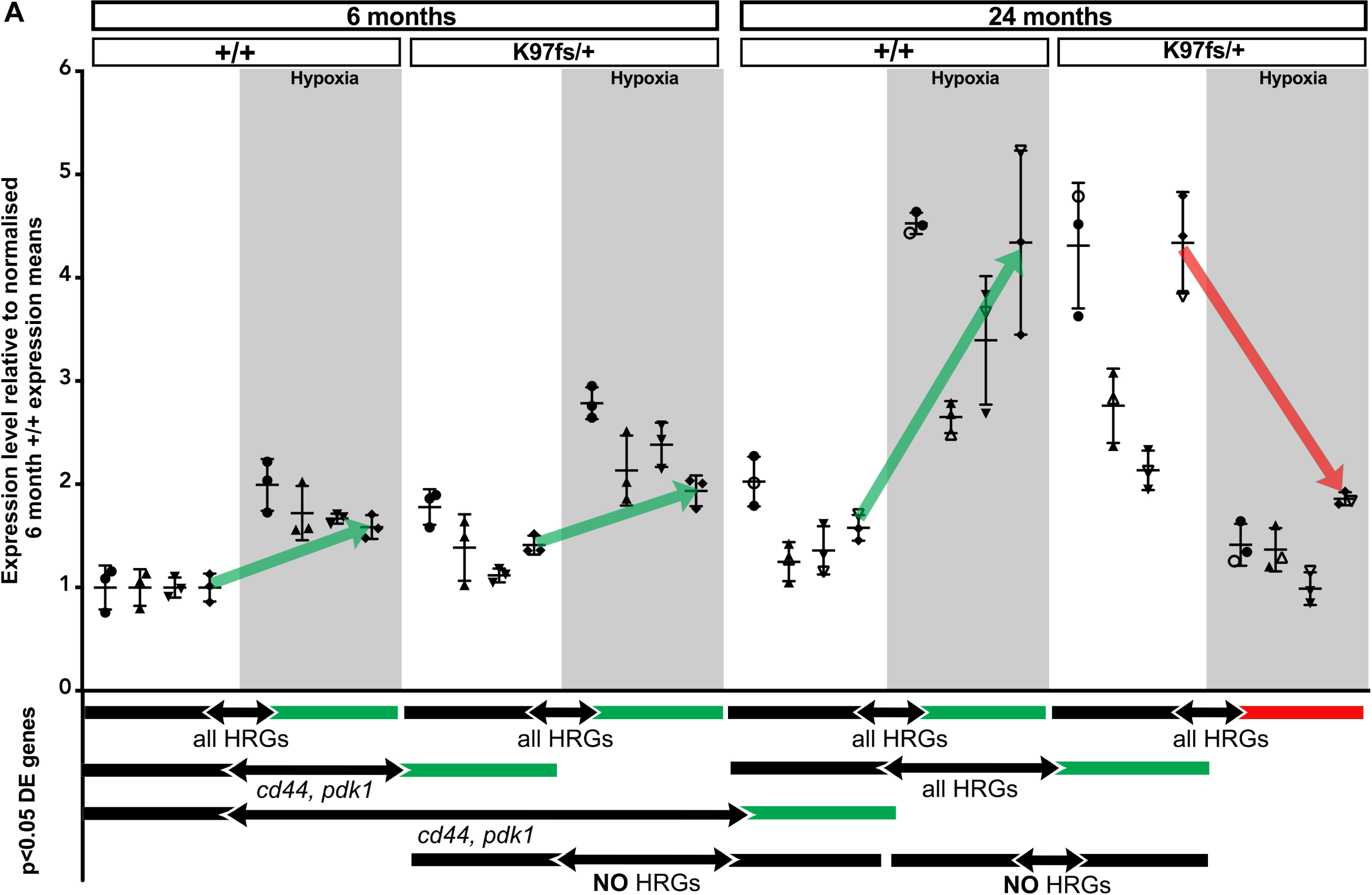
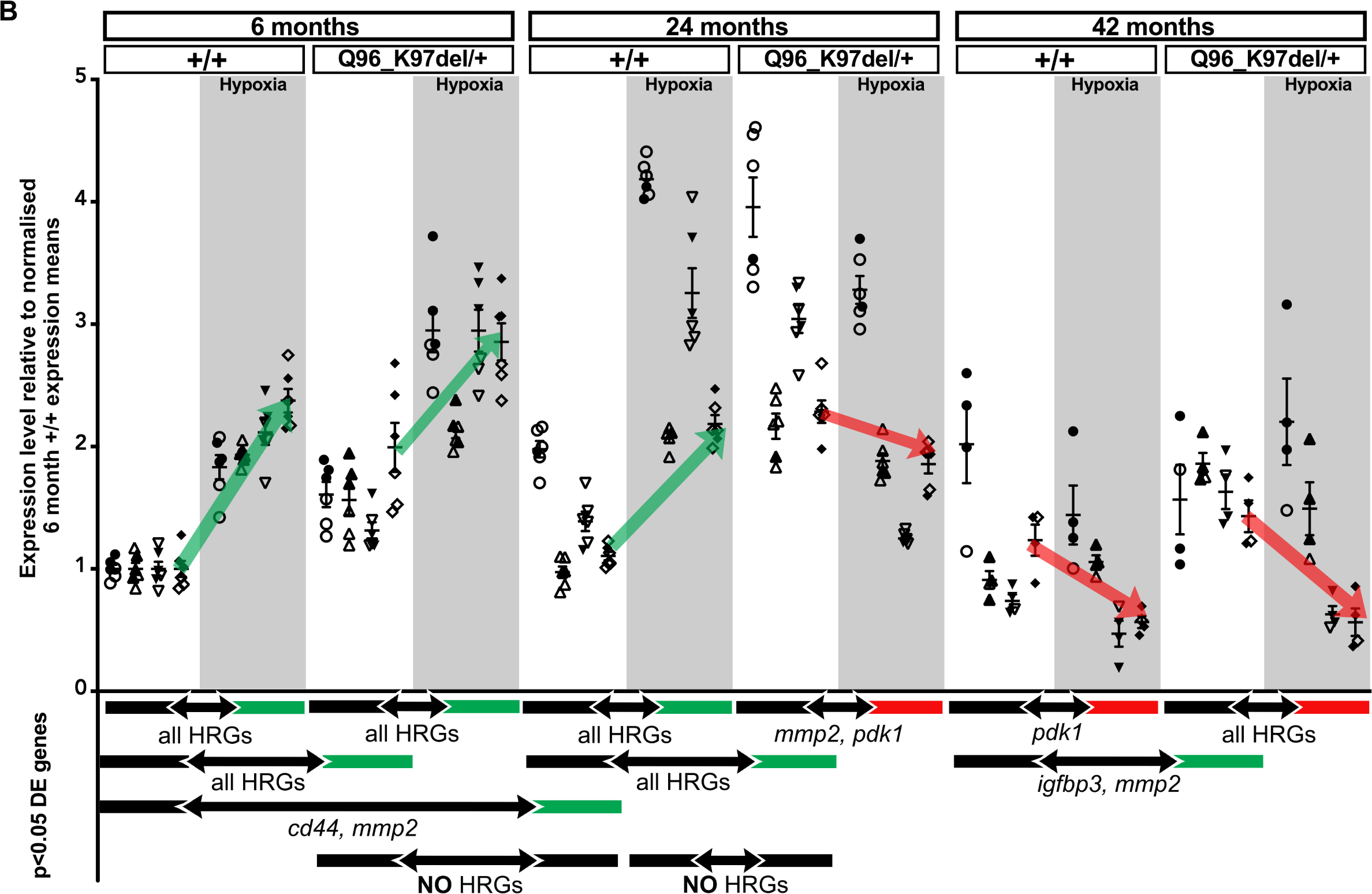
Hypoxia response gene expression in 6-month-old, 24-month-old and 42-month-old zebrafish brains under normoxia and hypoxia. Each data point on the graph indicates the relative transcript level for *cd44* (●), *igfbp3* (▴), *mmp2* (▾) and *pdk1* (♦) in 50ng of cDNA generated from a single zebrafish brain RNA sample, assuming reverse transcription is complete (filled symbols: females, outlined symbols: males). Copy numbers for each transcript in each sample were scaled to the normalized means of the transcript copy numbers in the wild type 6 months normoxia samples. The age and genotype of each sample is indicated at the top of the graph. Grey backgrounds indicate the hypoxia-treated samples. Wild type (+/+), *psen1*^K97fs/+^ (K97fs/+) heterozygous mutant and *psen1*^Q96_K97del/+^ (Q96_K97del/+) heterozygous mutant. p-value < 0.05 for differentially expressed (DE) genes determined by t-test: Two-Sample Assuming Unequal Variances. Solid lines flanking bi-directional arrows indicate comparisons in t-tests. The genes significantly differentially expressed are indicated below the bidirectional arrow. HRGs are HIF-1-responsive genes. The colour of the solid lines indicates the direction of differential expression; green indicates increased expression and red indicates decreased expression. Green and red arrows indicate the direction of *pdk1* expression change under hypoxia (arrow ends connect *pdk1* expression means). The raw dqPCR data, p-values and fold changes for all comparisons made are given in Tables 1 and 2 and Supplementary Data 1. **A.** Hypoxia response gene expression in K97fs/+ and +/+ siblings (6 months and 24 months). **B.** Hypoxia response gene expression in Q96_K97del/+ and +/+ siblings (6, 24 and 42 months).

The greatest surprise came when aged, 24-month-old brains heterozygous for either *psen1* mutation were subjected to acute hypoxia. Rather than further upregulating HRG expression, they either failed to do this or they even downregulated HRG expression (Figure 1A, B, p-values in Tables 1 and 2, Supplementary data 1). This inversion of the expected hypoxia response was particularly noticeable for expression of *pdk1*, suggesting an inability to upregulate anaerobic glycolysis in aged EOfAD-related mutant zebrafish under hypoxia. Assays of lactate content (a marker of anaerobic glycolysis) in brains both from young (7 months) and aged (31 months) *psen1*^*Q96_K97del*^ fish under acute hypoxia were consistent with this predicted failure to upregulate anaerobic glycolysis (Figure 2). However, in contrast to *pdk1* expression, there was no apparent upregulation of anaerobic glycolysis in young wild type brains under acute hypoxia (an observation also reported for young mouse brains under chronic hypoxia [43]).

**Figure 2.**
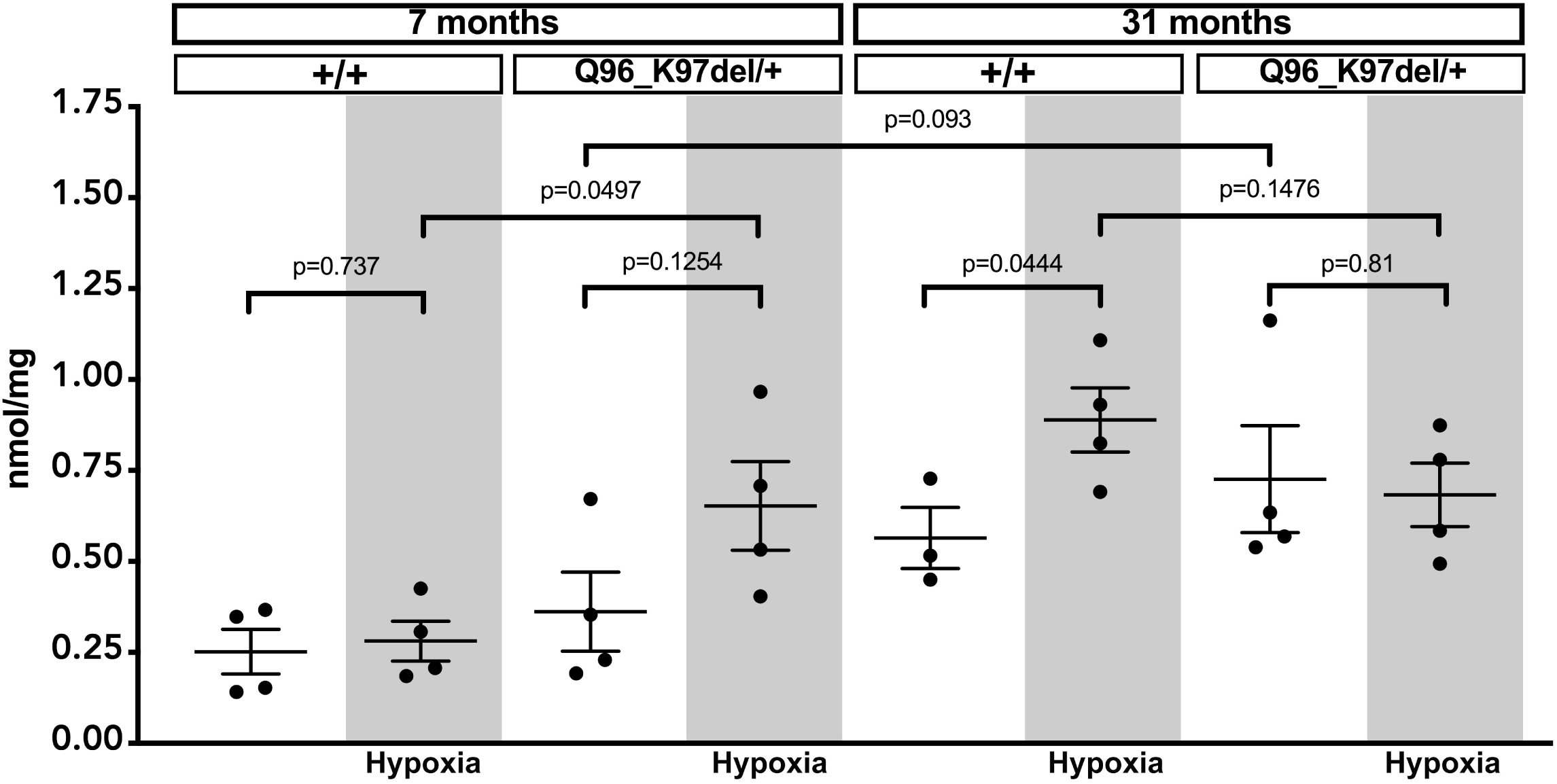
Lactate in 7-month-old and 31-month-old zebrafish brains under normoxia and hypoxia. Each data point on the graph indicates lactate content in a single whole zebrafish brain. All zebrafish were female. The age and genotype of each sample is indicated at the top of the graph. Grey backgrounds indicate the hypoxia-treated samples. Wild type (+/+) and *psen1*^Q96_K97del/+^ (Q96_K97del/+) heterozygous mutant. P-values are calculated using t-test: Two-Sample Assuming Unequal Variances. Raw lactate assay data is given in Supplementary Data 2. Lactate concentration is expressed as nmol per mg of whole brain from Q96_K97del/+ zebrafish and their +/+ siblings.

### Does decreased function of wild type psen1 accelerate brain aging?

Since age is the greatest risk factor for AD, the idea that EOfAD mutations of *psen1* might accelerate brain ageing seemed a parsimonious explanation for their pathogenicity. Therefore, we were surprised that both the less and the more EOfAD-like mutations of *psen1* appear to accelerate aging in terms of HRG responses. This suggests the possibility that the levels of wild type *psen1* allele expression are critical to the acceleration of brain aging rather than the presence of the mutations themselves. Unfortunately, despite repeated attempts, so far we have been unable to generate an antibody with which to assay zebrafish Psen1 protein levels. However, dqPCR allows us to assay and compare the relative levels of wild type and mutant *psen1* transcripts in zebrafish brains.

Premature termination codons commonly cause decreased mRNA transcript stability though the process of nonsense mediated decay (NMD, reviewed by [44]). The *psen1*^*K97fs*^ allele results in a premature termination codon (after 11 frameshifted amino acids) in exon 4 and its expression level is significantly decreased compared to transcripts from the wild type *psen1* allele [31] (displayed again in Figure 3A). This supports that the effects of *psen1*^*K97fs*^ are due to loss of function. Indeed the expression level of the wild type allele in the *psen1*^*K97fs*^ heterozygous fish brains is approximately half that of wild type sibling fish brains at both 6 months and 24 months of age [31] (displayed again in Figure 3A). There is no evidence of upregulation of the wild type allele in the *psen1*^*K97fs*^ heterozygous fish such as might be caused by the phenomenon of “genetic compensation” that can be driven by NMD [45]. Note however that, if the *psen1*^*K97fs*^ allele produces a truncated protein product, it is not expected to be completely without function (see Discussion).

**Figure 3.**
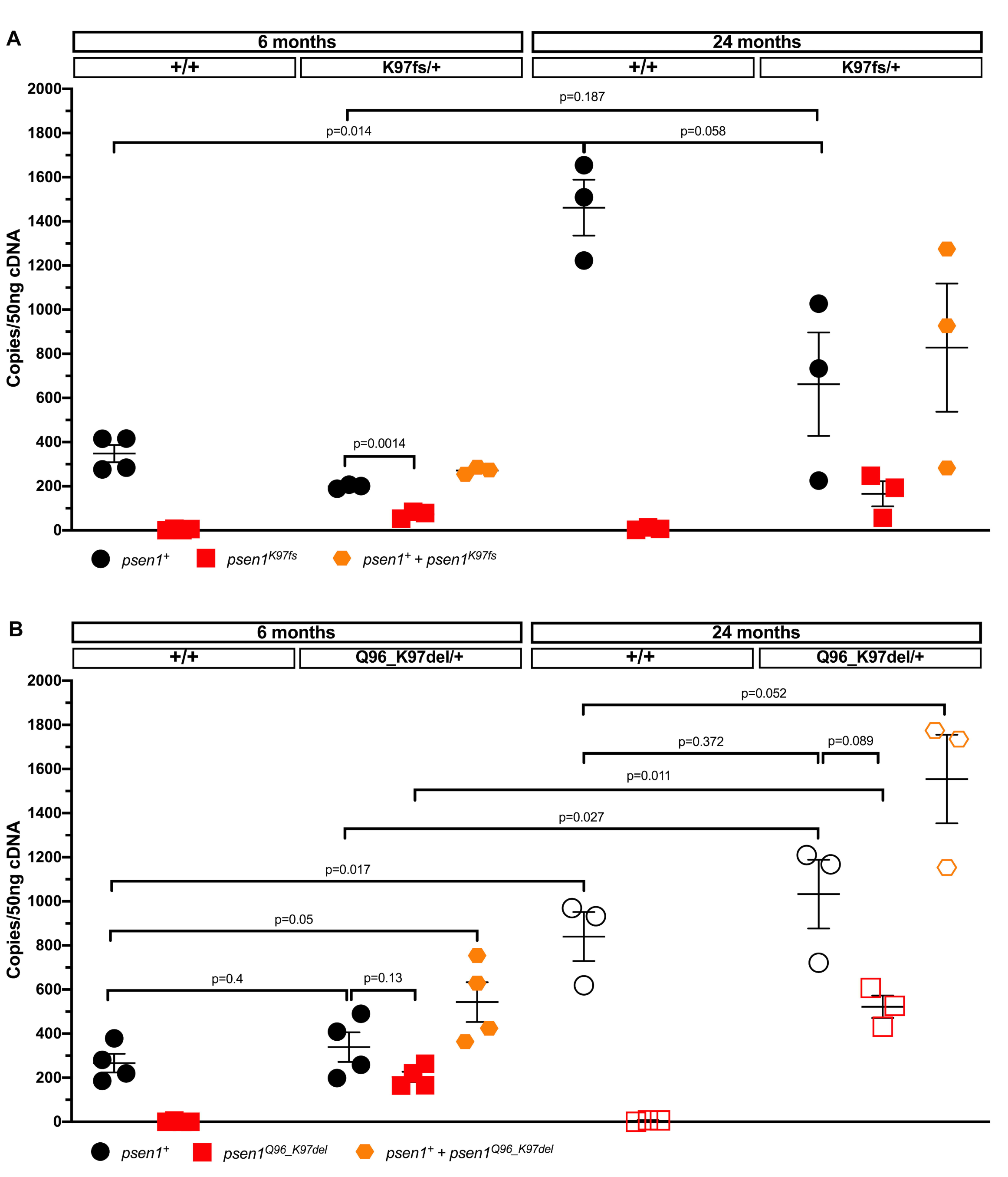
Allele-specific *psen1* transcript expression in 6-month-old and 24-month-old zebrafish brains. Each data point on the graph indicates *psen1* transcript copy number in 50ng of cDNA generated from a single zebrafish brain RNA sample, assuming reverse transcription is complete. The age and genotype of each sample is indicated at the top of the graph. Wild type (+/+), *psen1*^K97fs/+^ (K97fs/+) heterozygous mutant and *psen1*^Q96_K97del/+^ (Q96_K97del/+) heterozygous mutant. P-values are calculated using t-test: Two-Sample Assuming Unequal Variances. Raw dqPCR data and p-values are given in Supplementary Data 4. **A.** Allele-specific expression of either wild type *psen1* (+, black circles) or mutant *psen1* (K97fs, red squares) or *psen1*^*+*^ and *psen1*^*K97fs*^ combined (orange hexagons) (all females). **B.** Allele-specific expression of wild type *psen1* (+, black circles) and mutant *psen1* (Q96_K97del, red squares) or *psen1*^*+*^ and *psen1*^*Q96_K97del*^ combined (orange hexagons) (filled symbols: females, outlined symbols: males). The 6-month-old and 24-month-old +/+ data is also included in Figure 5. They are included/repeated in this figure for comparison to data from their heterozygous mutant siblings.

The mutant transcripts in the EOfAD-like *psen1*^*Q96_K97del*^ heterozygous fish brains possibly show somewhat decreased expression compared to wild type transcripts although this was not statistically significant (at the p<0.05 level, Figure 3B). However, unlike in the *psen1*^*K97fs*^ heterozygous fish, the level of wild type allele expression in heterozygous *psen1*^*Q96_K97del*^ fish brains is similar to that seen in wild type siblings of the same age. Something must drive this increased level of wild type allele expression. If the accelerated aging of these fish is due to decreased wild type allele function, then this could be due to interference from the mutant protein product of the *psen1*^*Q96_K97del*^ allele (see Discussion).

### Evidence for accelerated brain aging in human EOfAD

Mutations of human *PSEN1* (or *PSEN2*) that prevent any ORFs from utilising the original termination codon have never been seen to cause EOfAD (in line with the “reading frame preservation rule” [25]). A notable difference between the zebrafish *psen1*^*K97fs*^ and *psen1*^*Q96_K97del*^ alleles is that only the latter obeys the reading frame preservation rule and only this EOfAD-like mutation upregulates wild type *psen1* allele expression to equal or exceed that in wild type fish. Does a similar upregulation of *PSEN1* gene expression occur in human EOfAD? To test this we used dqPCR to examine wild type and mutant PSEN1 allele expression in three post-mortem EOfAD brains (medial temporal cortex) and in healthy, age-matched controls (Figure 4). Consistent with our *psen1*^*Q96_K97del*^ mutant data, transcripts of the mutant human mutant *PSEN1* allele was always detectable and at approximately half the level of the transcripts from the wild type allele. However, expression of the wild type allele was lower than in the age-matched controls (Figure 4). We reasoned that, since mutation of *psen1* in zebrafish appears to accelerate brain aging, and since the human EOfAD brains were from people who had died from this disease, their decreased human *PSEN1* expression might represent a phenomenon more similar to gene expression in extremely aged fish brains. Therefore, we compared the relative *psen1* gene expression levels between the young (6 months), aged (24 months) and extreme aged (51 months) wild type female fish brains. (Mutant fish brain cDNAs were not available for this experiment.) We observed the expected increase in *psen1* expression from 6 to 24 months but then a significant decrease in expression by 51 months (Figure 5). This supports that human brains with overt EOfAD may display the equivalent of an “extremely aged” molecular state giving them a lower level of *PSEN1* expression than chronologically age-matched healthy brains that are, in reality, less aged biologically.

**Figure 4.**
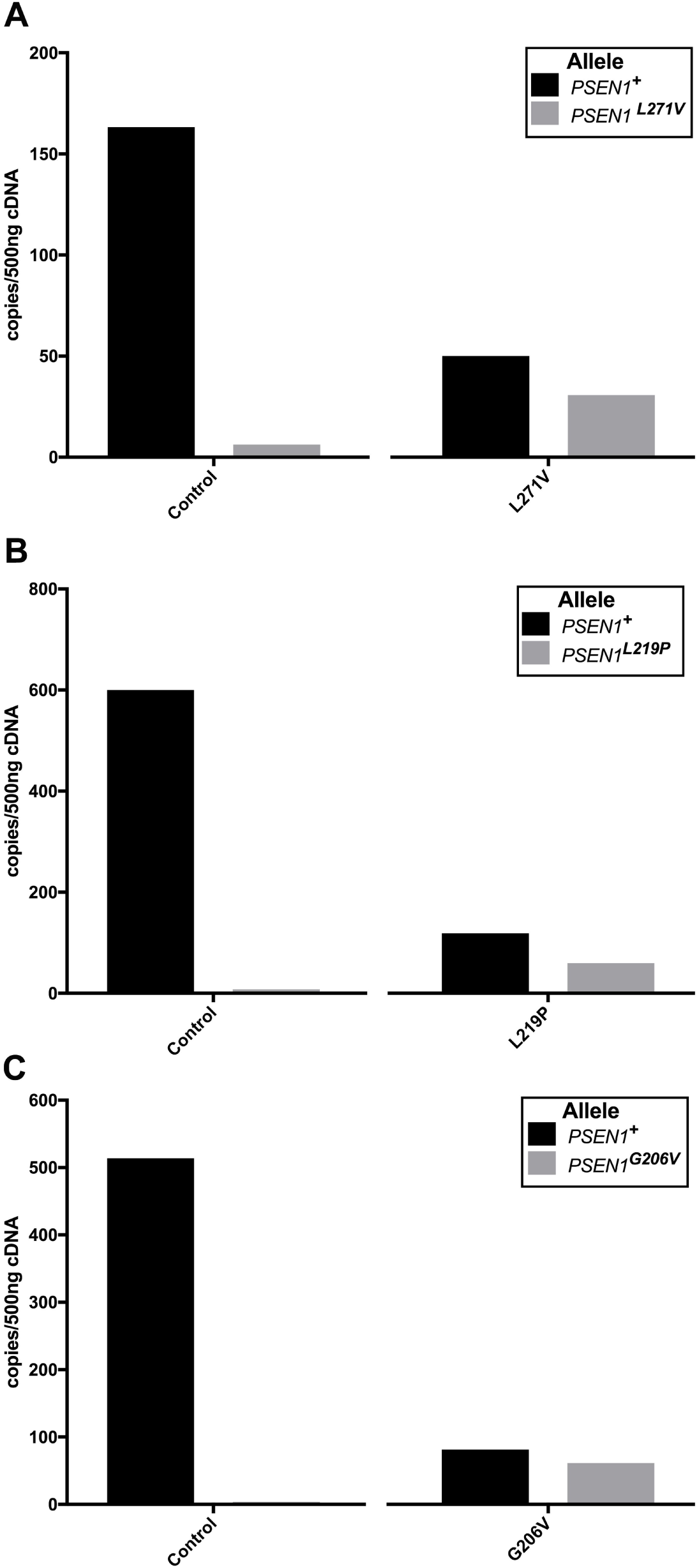
*PSEN1* transcript expression in the middle temporal gyrus region of brains of *PSEN1* mutation carriers and age-matched controls. Age-matched control (control, with corresponding mutation carrier sample in parentheses), *PSEN1*^L271V/+^ heterozygous mutation carrier (L271V/+), *PSEN1*^L219P/+^ heterozygous mutation carrier (L219P /+) and *PSEN1*^G206V/+^ heterozygous mutation carrier (G206V/+). Raw dqPCR data is given in Supplementary Data 3. **A.** Expression of wild type (*PSEN1*^+^) and L271V mutant (*PSEN1*^L271V^) alleles in control (51 years old) and L271V/+ (51 years old) cDNA samples. **B.** Expression of either wild type (*PSEN1*^+^) and L219P mutant (*PSEN1*^L219P^) alleles in control (60 years old) and L219P/+ (61 years old) cDNA samples. **C.** Expression of either wild type (*PSEN1*^+^) and G206V (*PSEN1*^G206V^) alleles in control (29 years old) and G206V/+ (32 years old) cDNA samples. The normalised *PSEN1* gene expression data (two different reference genes, *CYC1* and *RPL13*) is given in Supplementary Data 3.

**Figure 5.**
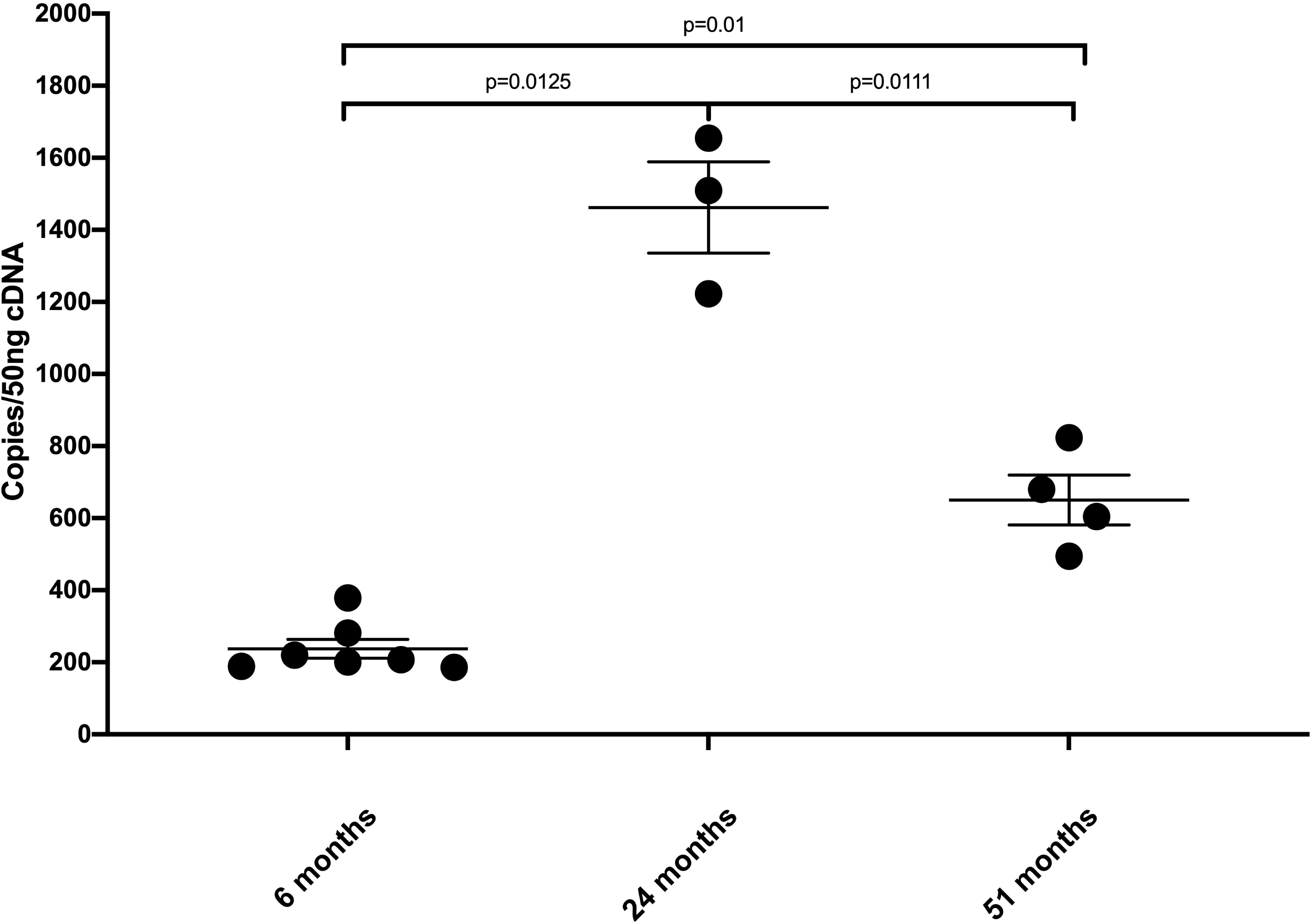
Wild-type *psen1* allele expression in 6-month-old, 24-month-old and 51-month-old wild-type zebrafish brains. Each data point on the graph indicates *psen1* transcript copy number in 50ng of cDNA generated from a single zebrafish brain RNA sample, assuming reverse transcription is complete. All samples are wild type females. The age of each sample is indicated at the bottom of the graph. P-values are calculated using t-test: Two-Sample Assuming Unequal Variances. Raw dqPCR data is given in Supplementary Data 3.

Interestingly, the age-dependent changes in brain transcript expression of a zebrafish paralogue of *HIF1α* (*hif1ab*) mirror closely those of *psen1* expression but are very different from its protein expression (see below and Figure 6).

**Figure 6.**
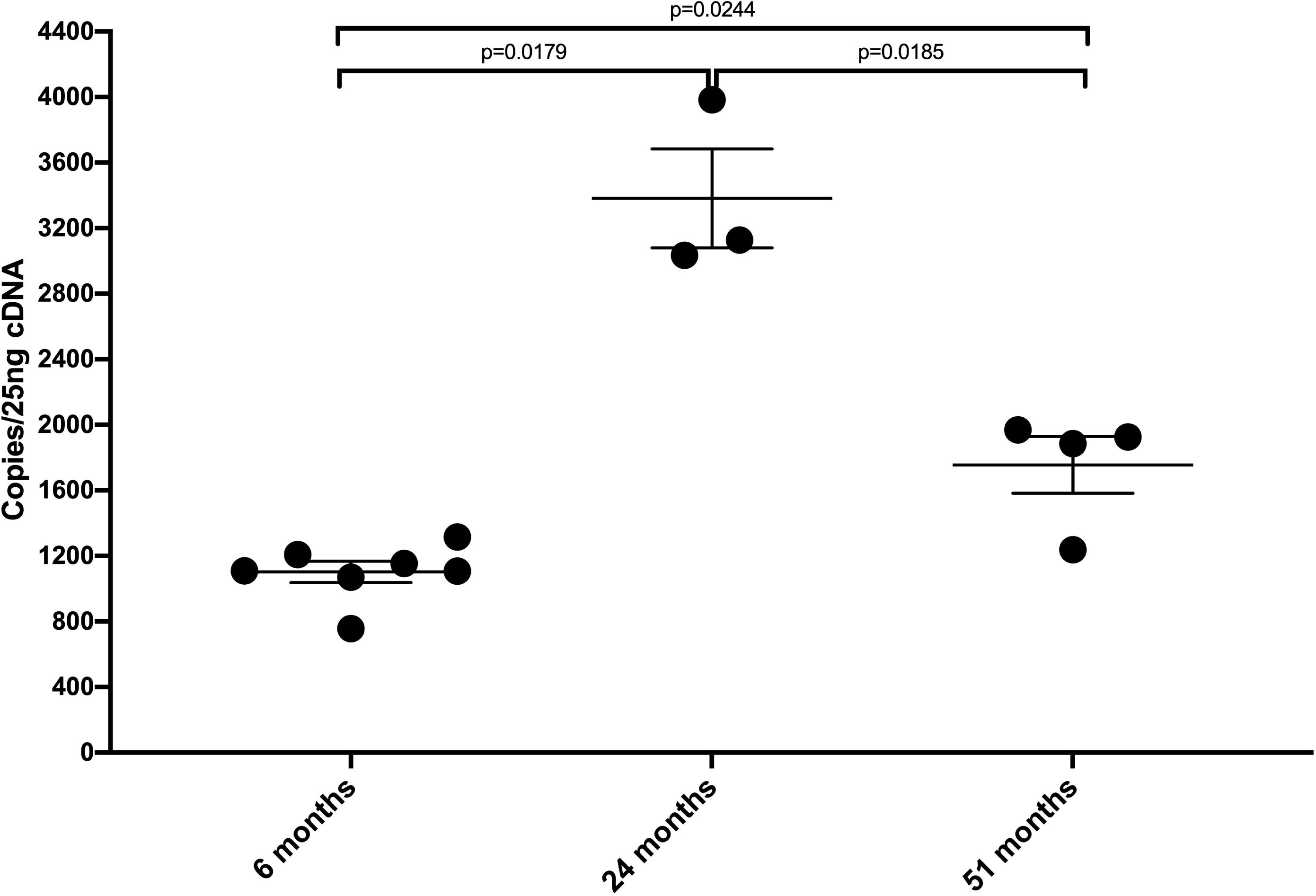
*Hif1ab* allele expression in 6-month-old, 24-month-old and 51-month-old wild-type zebrafish brains. Each data point on the graph indicates *psen1* transcript copy number in 25ng of cDNA generated from a single zebrafish brain RNA sample, assuming reverse transcription is complete. All samples are wild type female. The age of each sample is indicated at the bottom of the graph. P-values are calculated using t-test: Two-Sample Assuming Unequal Variances. Raw dqPCR data is given in Supplementary Data 4.

### Loss of HIF1 in aged brains is conserved across vertebrate species

HIF1α is one component of the heterodimeric transcription factor HIF1 that plays an important role in cellular responses to hypoxia. Hypoxia increases the stability of HIF1α protein, thus increasing HIF1 levels and upregulating a large number of HRGs. In a study of brain aging and responses to hypobaric hypoxia in rats, Ndubuizu et al [46, 47] observed an inability to stabilise Hif1α under hypoxia after 18 months of age. To test whether this phenomenon might explain the inability of extremely aged zebrafish brains to upregulate HRGs in response to acute hypoxia, we used an antibody against the paralogous zebrafish protein Hif1ab. We observed that wild type zebrafish already fail to stabilise Hif1ab from 24 months of age (Figure 7A and B), at a time when their basal HRG expression is raised compared to younger fish, and during which they can still greatly increase HRG expression under hypoxia (as shown in Figure 1). This shows that a failure to stabilise Hif1a in aged vertebrate brains is a characteristic conserved over an evolutionary period of greater than 430 million years, (i.e. since the divergence of the teleost and tetrapod lineages [48]) but this does not explain the “inversion” of HRG expression responses to hypoxia in aged zebrafish brains.

**Figure 7.**
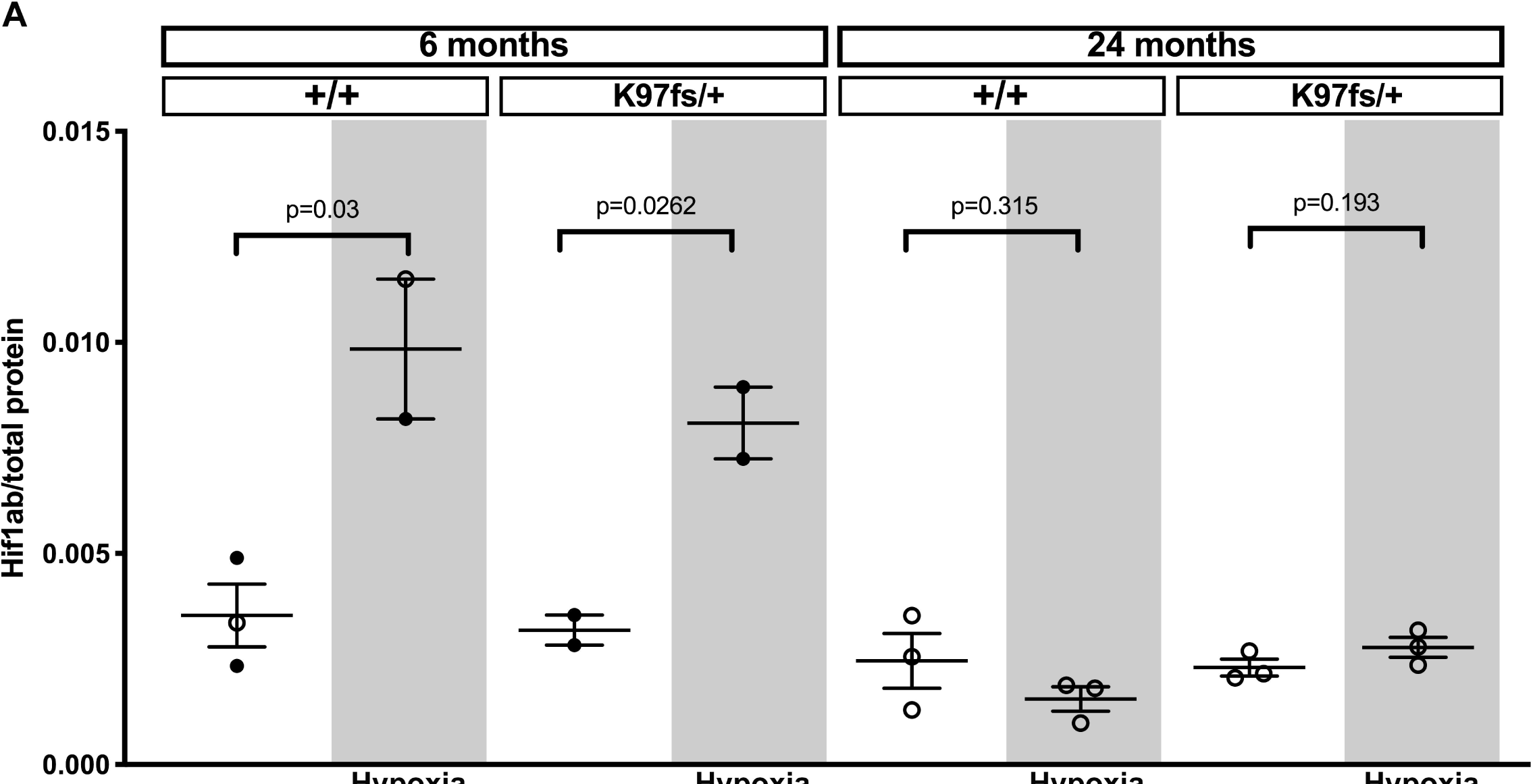
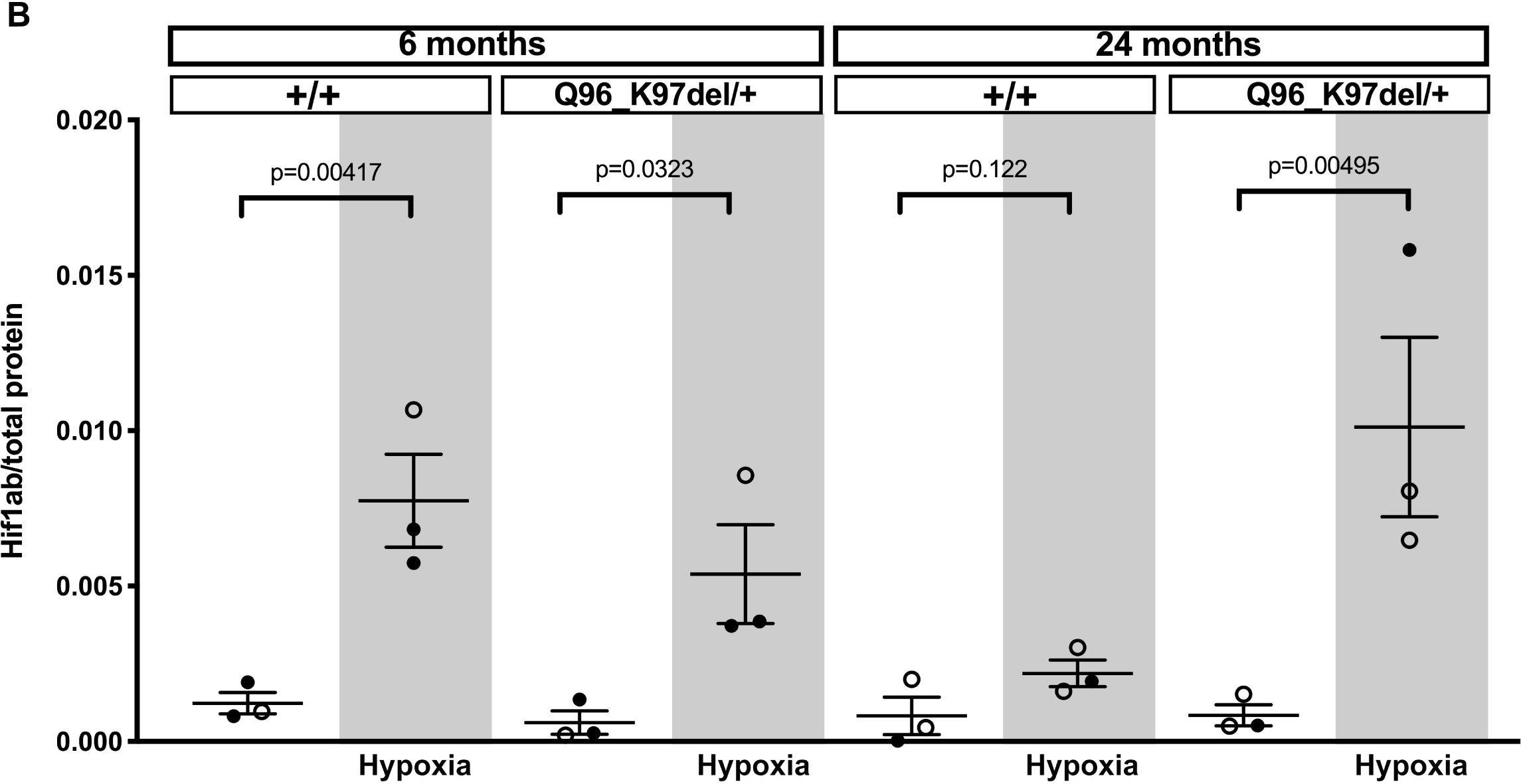
Hif1ab protein levels in 6-month-old and 24-month-old zebrafish brains under normoxia and hypoxia. Each data point on the graph indicates the Hif1ab protein level in protein extracted from a single zebrafish brain (filled circles: females, outlined circles: males). The age and genotype of each sample is indicated at the top of the graph. Grey backgrounds indicate the hypoxia-treated samples. Wild type (+/+), *psen1*^K97fs/+^ (K97fs/+) heterozygous mutant and *psen1*^Q96_K97del/+^ (Q96_K97del/+) heterozygous mutant. P-values are calculated using t-test: Two-Sample Assuming Unequal Variances on Ln transformed data. Raw densitometry data and p-values are given in Supplementary Data 5 and western blot images are shown in Supplementary Figure 2. **A.** Hif1ab protein level in K97fs/+ and +/+ siblings. **B.** Hif1ab protein level in Q96_K97del/+ and +/+ siblings.

### The EOfAD-like mutation of psen1 drives age-inappropriate stabilisation of a HIF1 paralogue under acute hypoxia

Only one previous study, De Gasperi et al. in 2010, has noted that PSEN1 protein directly physically interacts with HIF1α [37]. Using immortalised mouse embryonic fibroblasts these authors noted that Psen1 is required for stabilisation of Hif1α under chemical mimicry of hypoxia using CoCl_2_ or when treated with insulin. When mouse *Psen1* expression was replaced in these cells by human EOfAD mutant *PSEN1*^*M146V*^, stabilisation of Hif1α under chemical mimicry of hypoxia was normal but Hif1α stabilisation due to insulin treatment was deficient. Our analysis of Hif1ab stabilisation under acute hypoxia in aged, 24 month, zebrafish revealed a surprising difference between the heterozygous putative null mutant brains and EOfAD-like mutant brains. The EOfAD-like *psen1*^*Q96_K97del*^ brains stabilised Hif1ab under acute hypoxia, at an age when neither the wild type or *psen1*^*K97fs*^ brains did this (Figure 7). The consequences of this age-inappropriate stabilisation of Hif1ab are currently unknown but this observation emphasises the importance of observing EOfAD-like mutation activities in aged, non-transgenic animal models.

## Discussion

The production of energy supports all other cellular functions and is, therefore, fundamental to cellular, tissue, organ and organismal health. As the human brain ages, vascular function gradually declines [12, 13] making it more difficult to deliver oxygen and energy substrates to support the brain’s energy needs. As a consequence, transcription of *HIF1α* increases in the brain with age [49]. Increased HIF1, acting to increase expression of *PDK1*, would be expected to raise the rate of anaerobic glycolysis to maintain production of ATP (that is necessary, among other things, for maintaining the solubility of proteins to prevent the protein aggregation characteristic of neurodegenerative conditions [50]). However, a primary diagnostic marker of AD brains is that they are hypometabolic with both reduced oxygen and glucose consumption (as revealed by brain imaging studies, see review [12, 19]). Also, Liu *et al.* found evidence for reduced levels of HIF1α protein in post-mortem sporadic AD brains compared to normal, aged-matched controls [19]. Recently, machine learning was employed to analyse structural MRI data from healthy controls and patients with dementia and other neurological conditions [51]. This revealed distinct patterns of accelerated brain ageing in the dementia patients. These observations, together with our previous zebrafish *psen1* mutation data showing acceleration of ageing at the level of gene expression [31], and the hypoxia response data we have presented here, are consistent with a view of AD as both a consequence of ageing and as a pathological, hypometabolic brain state.

The clarity of the responses we have observed in whole zebrafish brains is due to our ability to place these entire, tiny ∼7 mg brains under acute hypoxia. The human brain is ∼250,000-fold larger in mass than a zebrafish brain, and the hypoxia it experiences as a result of failing vascular function would be expected to be regional (i.e. depending on the energy demand and vascularisation of different brain regions) and would spread progressively as vascular function decreases [52, 53]. Notably, Raz et al [54] recently observed that hypoxic hypoperfusion in rat brain drives hyperphosphorylation of tau while Mesulam [55] noted over two decades ago that the spread of neurofibrillary tangles of hyperphosphorylated tau in human brains tracks closely with the cognitive changes of AD progression (in contrast to Aβ plaque pathology). This is further evidence that hypoxia plays an important role in AD pathogenesis.

Our data from this study are also consistent with the “stress threshold change-of-state” model of AD progression we presented previously [26]. That model postulates aged brains “inverting” (in terms of numerous molecular characteristics) into a pathological state as they reach a homeostatic threshold such as a limit in their ability to cope with the increasing oxidative stress caused by age-related hypoxia. A number of “inversion” phenomena during AD pathogenesis have previously been noted by others. For example, in a transcriptome-based comparison of various brain regions in normal aged control, mild cognitive impairment (MCI) and AD post-mortem individuals, Berchtold et al. [56] saw upregulation in MCI of genes involved in many cellular processes including mitochondrial energy generation. Many of these genes were subsequently downregulated in AD relative to controls. Similar observations have been made for the onset of dementia in Down Syndrome individuals [57]. Also, Arnemann et al. [58] also saw widespread elevation of metabolic brain network correlations during healthy aging but reduction of such correlations in Alzheimer’s disease.

Digital, quantitative PCR (dqPCR) facilitates comparison of the levels of different transcripts by allowing comparison of PCRs based on different primer pairs. Using dqPCR we could observe significantly decreased levels of *psen1*^*K97fs*^ transcripts compared to wild type transcripts, presumably due to the mutant transcripts including a premature termination codon and so being subject to NMD. This supports a mainly hypomorphic phenotype for this allele. Any truncated protein product of *psen1*^*K97fs*^ in heterozygous brains would lack its own γ-secretase catalytic activity. However, it would likely resemble the truncated PSEN2 isoform PS2V [59, 60] and so might enhance the γ-secretase activity provided by the wild type allele, as well as inhibiting the cellular unfolded protein response [59, 61]. In contrast, transcripts of the *psen1*^*Q96_K97del*^ allele were not significantly less stable than wild type transcripts in heterozygous brains, and their presence stimulated increased expression of the single wild type allele. We speculate that this might be due to the mutant protein product of *psen1*^*Q96_K97del*^ interfering with the function of the wild type Psen1 protein, either through interference with the formation of Psen1 holoprotein multimers [25], or through dominant inhibition of γ-secretase activity [62]. If either of these actions caused increased oxidative stress (e.g. by interfering with mitochondrial function [35, 63]), this might upregulate the expression of *psen1* [14]. Notably, in wild type fish we observed an increase in *psen1* and *hif1ab* transcript levels with age that subsequently decreased as the fish became extremely aged, demonstrating once again the “inversion” of gene expression with extreme age.

In post-mortem human carriers of three different *PSEN1* EOfAD mutations, we observed severely decreased *PSEN1* transcript expression in middle temporal gyrus tissue relative to healthy controls. If loss of *PSEN1* expression is also widespread in LOsAD brains, then this phenomenon may be important in Alzheimer’s disease pathogenesis. However, the published expression data for *PSEN1* in human LOsAD brain tissue shows varying results depending on the brain region studied, the method of measurement, and which particular transcript variant was measured. For example, in LOsAD brains, a decrease in expression of *PSEN1* mRNA has been observed in the hippocampus (along with *PSEN2*) [64] and in the frontotemporal region [65] while an increase in expression of *PSEN1* mRNA has been measured in the temporal lobes [66, 67] and superior frontal gyrus [67]. Other studies reported no significant difference in *PSEN1* expression in LOsAD in the frontal cortex [68, 69], the entorhinal cortex, the auditory cortex (although decreased *PSEN2* was observed) or the hippocampus [70]. It is difficult to form any firm conclusions on *PSEN1* mRNA expression in LOsAD from these inconsistent findings. This should encourage more detailed analysis of *PSEN1* expression in both EOfAD and LOsAD brains in the future.

While our observations are consistent with mutations in *PSEN* genes causing accelerated aging, there are numerous studies that question the idea of AD being an inevitable consequence of age [71-73]. Even if the failure to upregulate glycolysis under hypoxia we observed in extremely aged zebrafish is related to the brain hypometabolism of AD, this does not show that AD is an inevitable consequence of aging. Rather, our results only support that age is a major risk factor for AD and may even be a necessary precondition for the disease.

Our study has uncovered a number of phenomena that we currently cannot explain. Why is it that, under acute hypoxia, two year old zebrafish brains cannot stabilise Hif1ab but are still able to upregulate HRGs? Are there HIF1-independent systems activating HRGs such as PGC1α (a key regulator of cellular energy metabolism) [47]? Why does heterozygosity for only the EOfAD-like mutation in *psen1* (that does not truncate the open reading frame) allow unexpected stabilisation of Hif1ab under hypoxia at an age when upregulation of HRGs fails? Indeed, why do transcript levels of HRGs decrease under hypoxia in these fish rather than increasing or, at least, not changing? Clearly, neither age nor the presence of mutations in *psen1* alone is sufficient to explain these phenomena and an interaction between the two factors is involved.

HIF1α has been shown to interact physically with PSEN1 protein [37, 38] and a physical interaction of HIF1α with the γ-secretase complex (presumably via PSEN1) can increase γ-secretase activity [38]. In turn, increased γ-secretase activity can increase HIF1α stability (and so HIF1 activity) via a p75NTR-dependent feed-forward mechanism [74]. For this reason, and the known stabilisation of HIF1α by ferrous iron deficiency [75, 76] (that can be caused by failure of lysosomal acidification [77]), we are unable to assert with certainty that the increased basal HRG expression seen in our heterozygous *psen1* mutant brains is due to hypoxia. Further work will be required to dissect the influences of *psen1* mutations on HRG expression in young brains and to understand the inappropriate stabilisation of Hif1ab in aged, EOfAD-like *psen1* mutant brains.

The experimental results described in this paper emphasise the intimate interaction between age, *PSEN1* (the majority locus for EOfAD mutations), and the HIF1 master controller of cellular hypoxia responses and energy metabolism. Maintaining vascular health and youthful brain hypoxia responses are likely key to avoiding or delaying the onset of AD. For human EOfAD mutation carriers, a recent systematic review has revealed an association between physical exercise and better cognitive outcomes at expected symptom onset ages [78]. This is consistent with physical exercise being one of the few known protective treatments against LOsAD [79-82].

## Methods

### Zebrafish husbandry and animal ethics

All experiments, except for digital PCRs for *psen1* transcript expression in 6-month-old brains, were performed using Tubingen-strain zebrafish maintained in a recirculated water system. This work with zebrafish was conducted under the auspices of the Animal Ethics Committee of the University of Adelaide (permit no. 31945) and Institutional Biosafety Committee (IBC Dealing ID 122210). The digital PCRs for *psen1* transcript expression in 6-month-old brain experiments were performed using samples from Tubingen and AB-strain zebrafish maintained in a recirculated water system. This work with zebrafish was conducted under the auspices of the Macquarie University Animal Ethics Committee (permit no. 2015/034).

### psen1 mutations

The isolation of the *psen*^*K97fs*^ and *psen1*^*Q96_K97del*^ mutations has previously been described [31, 83]. Mutations were only analysed in the heterozygous state in this study.

### Genomic DNA extraction and genotyping PCR of zebrafish tissue

DNA was extracted from adult zebrafish tail clips and genotyped via PCR as described in [31]. Primers used for genotyping PCR were synthesized by Sigma-Aldrich. Oligonucleotide sequences are given in Table 2.

### Hypoxia treatment of adult zebrafish

Male or female adult zebrafish at the desired age and genotype were treated in low oxygen levels by placing zebrafish in oxygen-depleted water for 3 hours (oxygen concentration of 6.6 ± 0.2 mg/L in normoxia and 0.6 ± 0.2 mg/L in hypoxia. Brains were subsequently used for digital PCR or western blotting analysis.

### Whole brain removal from adult zebrafish

Adult fish were euthanized by sudden immersion in an ice water slurry for at least 30 seconds before immediate decapitation. The entire brain was then removed from the cranium for immediate RNA or protein extraction. All fish brains were sampled during late morning/noon to minimise effects of circadian rhythms. For the dqPCR *psen1* allele-specific experiments on 6-months-old zebrafish, whole heads were stored in RNAlater (Ambion, Life Technologies) prior to brain removal and total RNA extraction.

### RNA extraction and cDNA synthesis of total adult zebrafish brain

Total RNA was extracted from whole zebrafish brain (∼10mg) using the QIAGEN RNeasy Mini Kit according to the manufacture’s protocol as described in [14, 59]. The RNA for dqPCR hypoxia response genes experiments, was DNase treated using RQ1 DNase from Promega according to the manufacturer’s instructions. The RNA for dqPCR *psen1* allele specific experiments, was DNase treated using the DNA-free™ Kit from Ambion, Life Technologies according to the manufacturer’s instructions. RNA was quantified using a Nanodrop spectrophotometer. cDNA was synthesized using random hexamers and the Superscript III First Strand Synthesis System (Thermo Fisher) according to the manufacturer’s instructions.

### RNA extraction and cDNA synthesis from human brain tissue

Human brain tissues were obtained from the Sydney Brain Bank at Neuroscience Research Australia and the NSW Brain Tissue Resource Centre at the University of Sydney. The brains were collected under institutional ethics approvals. Whole brains were sliced into blocks and fresh-frozen and stored at −80°C. Tissue samples from three cases with mutant *PSEN1* and three age- and gender-matched controls without significant neuropathology (see Table 1 for details) were processed for RNA extraction using TRIzol reagent (Invitrogen) following the manufacturer’s protocol. All procedures were carried out using RNase-free reagents and consumables. Five micrograms of RNA was reverse transcribed into cDNA using Moloney-murine leukemia virus reverse transcriptase and random primers (Promega, Madison, Wisconsin, USA) in a 20 μl reaction volume. RNA integrity was assessed with high resolution capillary electrophoresis (Agilent Technologies) and only RNA with RNA Integrity Number (RIN) value greater than 6.0 was used.

### 3D Quant Studio Digital PCR

Digital PCR was performed on a QuantStudio™ 3D Digital PCR System (Life Technologies). Reaction mixes contained 1X QuantStudio™3D digital PCR Master Mix, Sybr® dye, 200nM of specific primers and 12.5 - 500ng cDNA (500ng for *PSEN1* and *PSEN2*, 100ng for *CYC1* and *RPL13*, 50ng for *cd44a, igfbp3, psen1* 25ng for *pdk1* and 12.5 ng for *mmp2*). The reaction mixture (14.5μl) was loaded onto a QuantStudio™3D digital PCR 20 K chip using an automatic chip loader according to the manufacturer’s instructions. Loaded chips underwent thermo-cycling on the Gene Amp 9700 thermo-cycling system under the following conditions: 96°C for 10 min, 39 cycles of 60°C (59°C for *CYC1* and *PSEN1G206V*site) for 2 min and 98°C for 30 sec, followed by a final extension step at 60°C (or 59°C see above) for 2 min. The chips were imaged on a QuantStudio™ 3D instrument, which assesses raw data and calculates the estimated concentration of the nucleic acid sequence targeted by the Sybr® dye by Poisson distribution [84]. Primers used were synthesized by Sigma-Aldrich and are listed in Table 2.

### Protein extraction of total adult zebrafish brain

Each brain was homogenised in water supplemented with Sample Reducing Agent (10X, contains 500 mM dithiothreitol) (Thermo Fisher Scientific) and Complete Protease Inhibitors (Roche Life Sciences). LDS buffer (Thermo Fisher Scientific) was then added and the sample was briefly homogenised and placed at 90°C for 20 minutes. Each sample was divided into several aliquots. To generate a protein standard to allow quantitative comparison between signals on separate western blots, 8 male non-mutant adult zebrafish at 6 months of age were selected and placed under hypoxia as described in [14]. Protein was extracted from each brain as above. After extraction all homogenates were combined, thoroughly mixed and divided into several aliquots, with one aliquot being sufficient for one Nu-PAGE gel (i.e. 3×10ul for 3 lanes). All samples and standards were stored at −80°C until required.

### Protein Immunoblotting

Samples and standards were prepared by sonication for 10 minutes followed by incubation at 90°C for 5 minutes. PrecisionPlusProteinDualXtra ladder (Bio-Rad) (7μl), standards (3 x 10μl) and samples (6 x ∼18μl) were loaded on a 4-12% Bis-Tris Nu-PAGE (Life Technologies) gel for electrophoresis. The protein samples and standards were subsequently transferred to a PVDF membrane using the Mini Blot Module transfer system according to the manufacturer’s protocol (Thermo Fisher Scientific). To detect the Hif1ab protein, the membrane was initially blocked in 3%w/v skim milk powder followed by incubation in a 1/3000 dilution of the Hif1ab antibody (Gene Tex, cat no. GTX131826). The membrane was washed and incubated in a 1/2500 dilution of the rabbit-HRP secondary antibody (Sigma). After washing, Hif1ab protein was detected on the membrane using ECL detection reagents (Thermo Fisher Scientific) and visualised using the Chemi Doc Imaging System (Bio-Rad). The Hif1ab protein band can be observed at ∼100kDa. Using Image Lab software (Bio-Rad), densitometry analysis was performed for the Hif1ab protein band for each sample and standard. An average value was obtained for the standards for each membrane. Each sample value was then normalised to the average standard value.

Each protein sample was quantified using the EZQ protein quantification kit (Thermo Fisher Scientific) according to the manufacturer’s protocol. The standard normalised value for each sample was then expressed as Hif1ab per total protein loaded. Each membrane displayed 3 normoxia and 3 hypoxia samples from the same age (6 or 24 months) and the same genotype (+/+, *psen1*^*Q96_K97del*^/+ or *psen1*^*K97fs*^/+) together with the protein standard in 3 separate lanes.

### ^L-^Lactate Content Analysis of zebrafish brain

^L-^Lactate content was analysed using the Lactate Colorimetric Kit II (BioVision). Brains were removed from zebrafish adults in cold 1xPBS, then weighed and homogenized in the Lactate Assay Buffer provided in the kit. Each sample was then filtered through a 10kDa MW spin filter (Sartorius Stedium Biotech) to remove all proteins. The lactate content of the eluate from each zebrafish brain was determined using the kit according to the manufacturer’s protocol and the original brain weight was used to calculate the nmol of lactate per mg of brain tissue.

## Acknowledgements

This research was supported by a grant from Australia’s National Health and Medical Research Council (NHMRC), (#1126422). MN was also generously supported by a grant from the family of Lindsay Carthew.

Brain tissues were received from the NSW Brain Tissue Resource Centre and Sydney Brain Bank. These brain banks are supported by the NHMRC of Australia, The University of New South Wales, Neuroscience Research Australia, and the National Institute of Alcohol Abuse and Alcoholism (NIH (NIAAA) R24AA012725). G.M.H. is a National Health and Medical Research Council of Australia Senior Principal Research Fellow (#1079679). Tissues were processed in the Dementia and Movement Disorders Laboratory supported by Forefront, a collaborative research group dedicated to the study of non-Alzheimer disease degenerative disorders, funded by NHMRC grants (#1037746 and #1095127).

## Conflicts of Interests

Nothing to disclose.

## Supplementary Information

**Supplementary Figure 1. PCA of all genes expressed in whole brain samples.** The number of differentially expressed (DE) genes observed between male and female brains at 6 months: none; at 24 months: 19. The number of DE genes observed between *+/+* and Q96_K97del*/+* brains at 6 months: 278; at 24 months: 543. The distribution of DE genes in these comparisons supports that age and the presence of the *psen1* mutation appears to affect brain gene expression while the sex of the zebrafish does not influence greatly overall brain differential gene expression.

**Supplementary Figure 2. Hif1ab western blot images.**

**Supplementary Data 1. Hypoxia Response Genes dqPCR data.**

**Supplementary Data 2. Lactate colorimetric assay data.**

**Supplementary Data 3. *Psen1, PSEN1, CYC1* and *RPL13* dqPCR data.**

**Supplementary Data 4. Hif1ab dqPCR data.**

**Supplementary Data 5. Hif1ab western blot densitometry data.**

## Notes

### Competing Interest Statement

The authors have declared no competing interest.

### Summary of Updates

This new submission has been adjusted with updated results from experiments that were suggested from reviewers. We performed additional experiments of outcrossing the mutant zebrafish to another strain and repeated digital PCRs measuring psen1 allele expression. A different person performed these experiments and discovered that there were issues with the original experimental design, therefore the results in the initial submission of "biased psen1 allele expression" was an artefact. These results have been removed and replaced.

