## Supplementary figures and images for "Accelerated loss of hypoxia response in zebrafish with familial Alzheimer’s disease-like mutation of Presenilin 1"

### Supplementary Figure 2

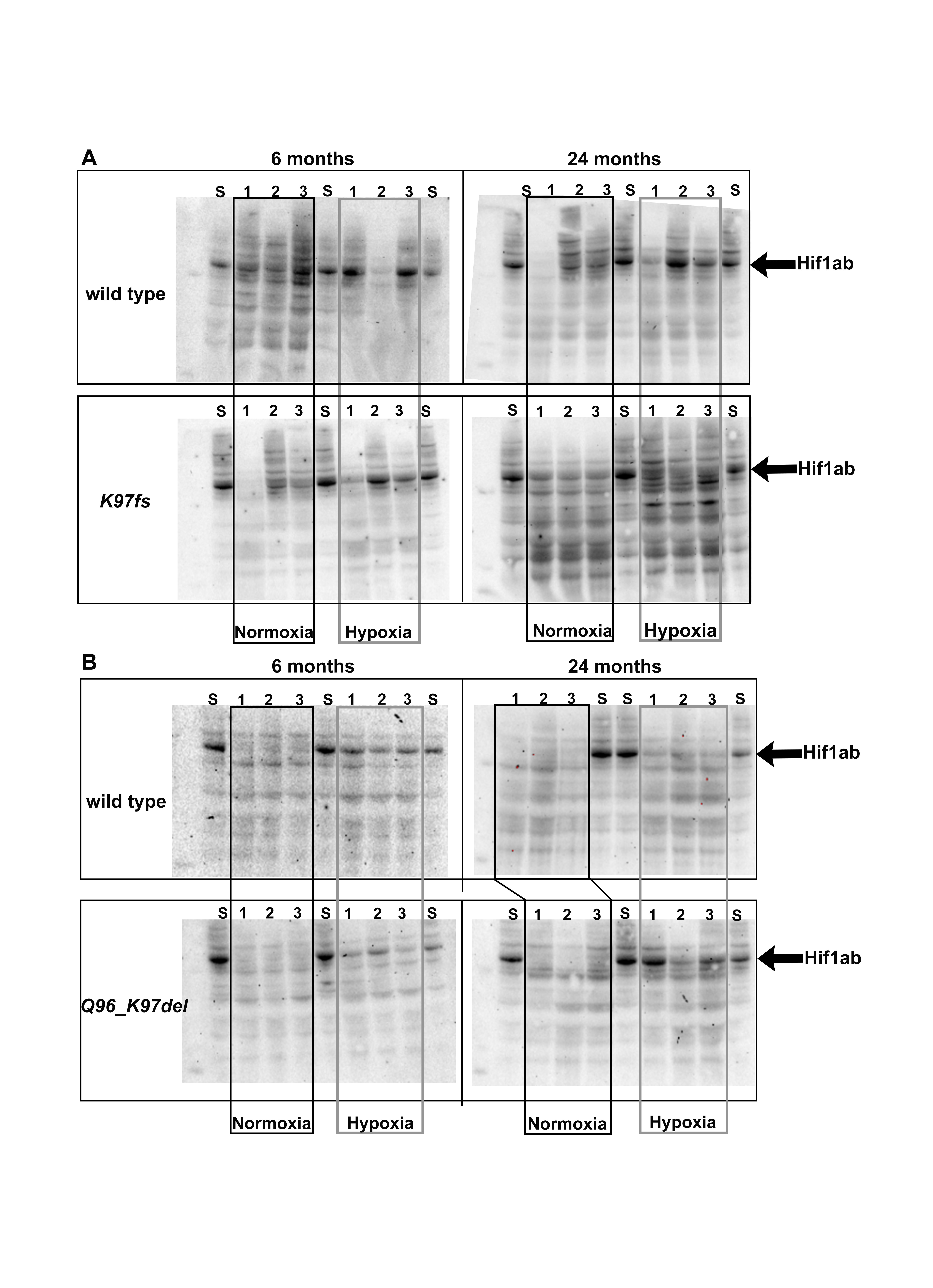

### Supplementary Figure 2

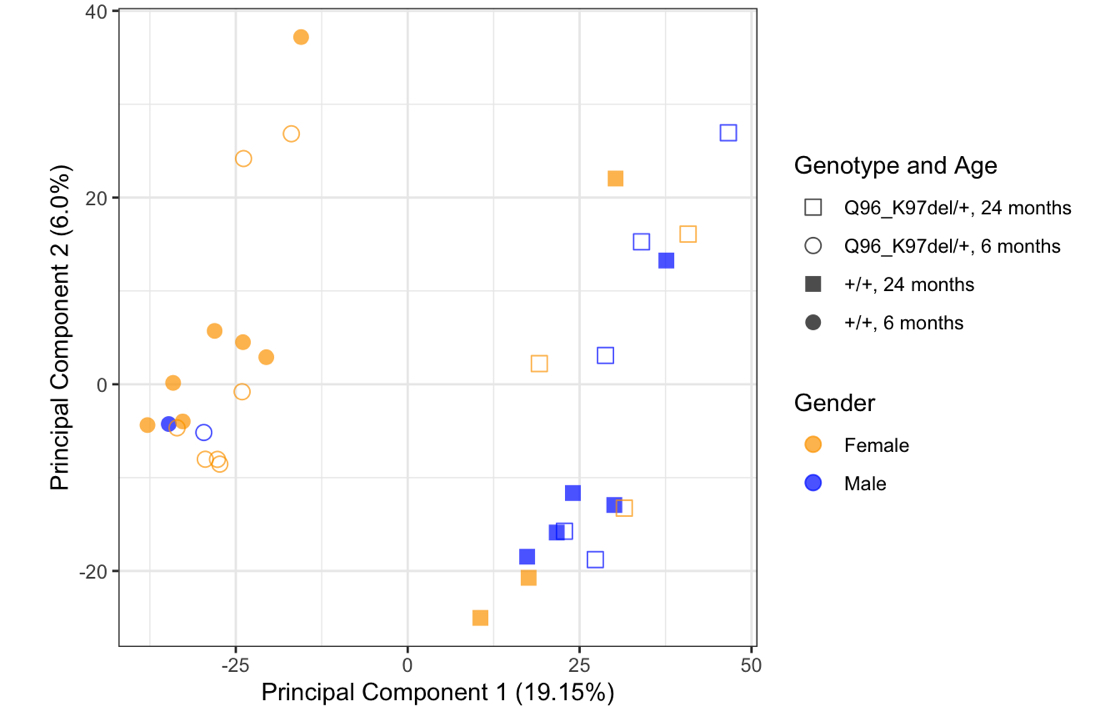
